# Msh2-Msh3 DNA-binding is not sufficient to promote trinucleotide repeat expansions in *Saccharomyces cerevisiae*

**DOI:** 10.1101/2024.08.08.607243

**Authors:** Katherine M. Casazza, Gregory M. Williams, Lauren Johengen, Gavin Twoey, Jennifer A. Surtees

## Abstract

Mismatch repair (MMR) is a highly conserved DNA repair pathway that recognizes mispairs that occur spontaneously during DNA replication and coordinates their repair. In *Saccharomyces cerevisiae*, Msh2-Msh3 and Msh2-Msh6 initiate MMR by recognizing and binding insertion deletion loops (in/dels) up to ∼ 17 nucleotides (nt.) and base-base mispairs, respectively; the two complexes have overlapping specificity for small (1-2 nt.) in/dels. The DNA-binding specificity for the two complexes resides in their respective mispair binding domains (MBDs) and have distinct DNA-binding modes. Msh2-Msh3 also plays a role in promoting *CAG/CTG* trinucleotide repeat (TNR) expansions, which underlie many neurodegenerative diseases such as Huntington’s Disease and Myotonic Dystrophy Type 1. Models for Msh2-Msh3’s role in promoting TNR tracts expansion have invoked its specific DNA-binding activity and predict that the TNR structure alters its DNA binding and downstream activities to block repair. Using a chimeric Msh complex that replaces the MBD of Msh6 with the Msh3 MBD, we demonstrate that Msh2-Msh3 DNA-binding activity is not sufficient to promote TNR expansions. We propose a model for Msh2-Msh3-mediated TNR expansions that requires a fully functional Msh2-Msh3 including DNA binding, coordinated ATP binding and hydrolysis activities and interactions with Mlh complexes that are analogous to those required for MMR.

**Article Summary:** The mismatch repair (MMR) protein complex Msh2-Msh3 promotes trinucleotide repeat (TNR) expansions that can lead to neurodegenerative diseases, while the Msh2-Msh6 complex does not. We tested the hypothesis that Msh2-Msh3’s specific DNA binding activity is sufficient to promote TNR expansions, using a chimeric MSH complex *in vivo* and *in vitro*. We found that the Msh2-Msh3-like DNA-binding was not sufficient to promote TNR expansions. Our findings indicate that Msh2-Msh3 plays an active, pathogenic role in promoting TNR expansions beyond simply binding to TNR structures.

## Introduction

Mismatch repair (MMR) is an evolutionarily conserved pathway that recognizes and corrects errors in DNA replication. Two heterodimeric MutS homolog (Msh) complexes initiate MMR through the recognition of distinct DNA structures that arise as a result of nucleotide misincorporation, leading to mispairs, or DNA polymerase slippage events, leading to insertions or deletions (in/dels). Msh2-Msh6, or MutSα, recognizes and binds mispairs and small in/dels (1-2 nucleotides (nt.)) through a conserved Phe-X-Glu motif that intercalates with the mispair, burying it within Msh2-Msh6 (Obmolova *et al*. 2000; Warren *et al*. 2007). Msh2-Msh3, or MutSβ and the focus of this study, recognizes and binds in/dels of up to 17 nucleotides, as well as some mispairs, through a conserved Tyr-Lys pair that interacts with the 5’ double-strand/single-strand DNA junction in loop structures, leaving at least part of the DNA structure accessible (Sia *et al*. 1997; Kunkel and Erie 2005; Lee *et al*. 2007; Dowen *et al*. 2010; Gupta *et al*. 2011).

The mispair binding domains (MBD) of Msh3 and Msh6 are responsible for Msh2-Msh3 versus Msh2-Msh6 structure-specific DNA binding activities and contain the highly conserved Phe-X-Glu (Msh6) or Tyr-Lys (Msh3) motifs, as well as other highly conserved residues that contribute to DNA structure specificity (Obmolova *et al*. 2000; Warren *et al*. 2007; Gupta *et al*. 2011). Replacing the Msh6 MBD with the Msh3 MBD in the context of Msh2-Msh6 (**Figure 1**) swapped the structure-binding specificity of the resulting Msh2-msh6(3MBD) complex, which exhibited a preference for Msh2-Msh3 in/del loop (IDL) substrates (Shell *et al*. 2007; Brown *et al*. 2016). The ATPase activity of Msh2-msh6(3MBD) was stimulated by Msh2-Msh3 substrates and this chimeric complex gained Msh2-Msh3’s ability to bypass protein blocks, “hopping” over nucleosomes (Brown *et al*. 2016), which Msh2-Msh6 lacks (Gorman *et al*. 2007; Brown *et al*. 2016). These data indicate that Msh2-Msh3’s DNA-binding specificity is largely mediated through its MBD.

**Figure 1.**
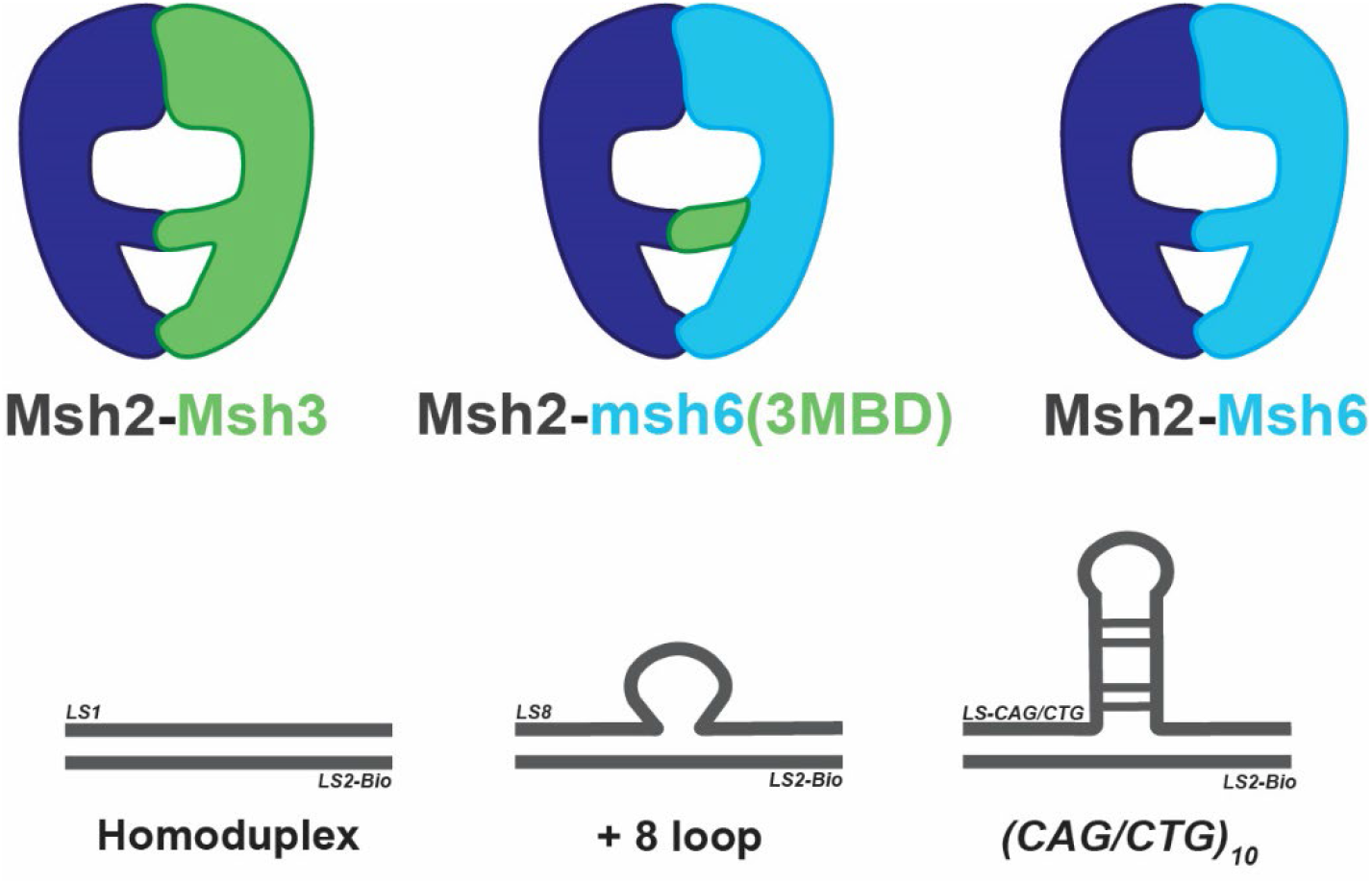
Schematic of MSH complexes and Msh2-Msh3 DNA substrates. *Top:* Schematic indicating the overall structure of the Msh2-Msh3, Msh2-Msh6 and chimeric MSH complexes. Msh2-msh6(3MBD) results from replacing the mispair binding domain (MBD) of Msh6 with that of Msh3, which is sufficient to switch DNA-binding specificities and DNA hopping capacity (Shell *et al*. 2007; Brown *et al*. 2016). *Bottom:* Schematic depicting the DNA structures used in this study: non-specific homoduplex DNA, an MMR-specific loop structure with 8 extrahelical nucleotides ([GT]_4_) and TNR tract slipped structures for *CAG* and *CTG* tracts. The DNA sequences are in **Table 1**.

Once bound, Msh complexes bind ATP and recruit MutL homolog (Mlh) heterodimeric complexes MutLα and/or MutLγ in an ATP-dependent manner. The latent endonuclease activity of the Mlh complexes is activated by Msh and PCNA, nicking the nascent DNA strand (Furman *et al*. 2021; Pannafino and Alani 2021). This is followed by recruitment of Exo1 and other downstream factors that promote excision, and resynthesis of the nascent DNA to complete repair (Jiricny 2006; Li 2008; Goellner *et al*. 2015; Keogh *et al*. 2017). ATP hydrolysis by Msh2-Msh3 is thought to be important for turnover of the complex, allowing it to re-bind DNA (Kijas *et al*. 2003; Owen *et al*. 2005; Owen *et al*. 2009; Kumar *et al*. 2014).

**Table 1:**
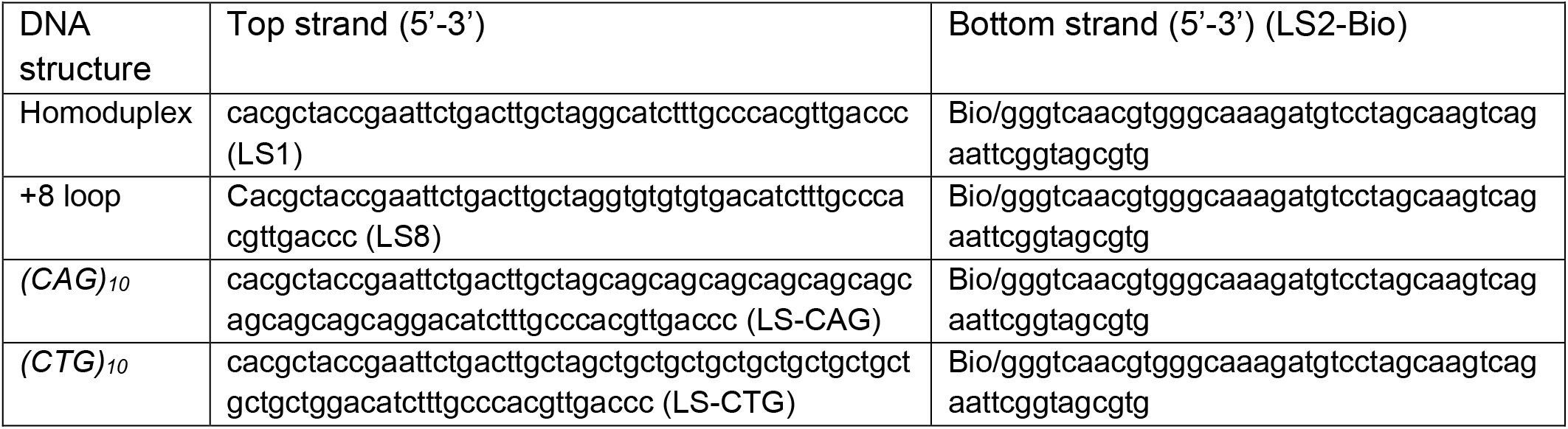
DNA substrates.

Loss of MSH3 compromises genome stability and leads to an increase in microsatellite instability (Marsischky *et al*. 1996; Sia *et al*. 1997; Harrington and Kolodner 2007; Lee *et al*. 2007; Kumar *et al*. 2013). Single nucleotide polymorphisms (SNPs) in human *MSH3* have been associated with a predisposition to cancer (Miao *et al*. 2015; DE’ ANGELIS *et al*. 2018; Santos *et al*. 2018; Aelvoet *et al*. 2023; Rashid *et al*. 2024). These genome-protective characteristics of *MSH3* are in direct contrast to its pathogenic role in promoting trinucleotide repeat (TNR) expansions (Kantartzis *et al*. 2012; Williams and Surtees 2015; Keogh *et al*. 2017). TNR expansions are the cause of over 40 neurodegenerative and neuromuscular diseases such as Huntington’s Disease and Myotonic Dystrophy type 1, which are caused by *CAG* and *CTG* expansions respectively (Mirkin 2007). *MSH3* promotes TNR expansions, including *CNG* tracts, in multiple model systems including yeast, mice, and human cell culture (Kantartzis *et al*. 2012; Lahue 2020; Iyer and Pluciennik 2021; Richard 2021; MATOS-RODRIGUES *et al*. 2023). In genome-wide association studies (GWAS), *Msh*3 was identified as a genetic modifier of TNR expansions; polymorphisms identified in *Msh3* that correlated with higher *Msh3* expression, exhibited increased TNR tract instability in mice (TOMÉ *et al*. 2013; Bettencourt *et al*. 2016; Lee *et al*. 2019). In contrast, *MSH6* does not promote significant *CNG* expansions (Kantartzis *et al*. 2012; Iyer and Pluciennik 2021), indicating that MMR-dependent TNR expansions are specific to the Msh2-Msh3 mediated pathway, leading us to focus on the molecular activities of Msh2-Msh3 that are necessary for promoting TNR expansion.

Msh2-Msh3 recognizes and binds the slipped strand secondary structures thought to form during replication of TNR tracts, with affinities similar to in/dels (Owen *et al*. 2005; Tian *et al*. 2009; Lang *et al*. 2011). However, the execution of repair is altered, leading to TNR tract expansions rather than repair pathways that maintain tract length. While Msh2-Msh3 ATPase activity is likely required for TNR expansions (TomÉ *et al*. 2009; Keogh *et al*. 2017), TNR DNA structures decreased Msh2-Msh3 ATP-binding and hydrolysis activities compared to that observed with an in/del MMR substrate (Owen *et al*. 2005; Owen *et al*. 2009). Similarly, Msh2-Msh3 nucleotide-binding and hydrolysis activities are differentially modified in the presence of DNA structures it binds in double-strand break repair versus MMR (Kumar *et al*. 2014). These observations led to the prediction that Msh2-Msh3 becomes “trapped” upon binding TNR structures, preventing repair of these slipped-strand secondary structures (Owen *et al*. 2005; Lang *et al*. 2011). Our *in vitro* work has suggested that at least part of Msh2-Msh3’s role in promoting TNR expansions is in stabilizing the TNR structures (Kantartzis *et al*. 2012), consistent with other studies (Tian *et al*. 2009). In this study, we sought to determine whether Msh2-Msh3’s specific DNA binding activity, through the Msh3 MBD, is sufficient to promote TNR expansions.

## Materials and Methods

### Protein Purification

Msh2-Msh3 was overexpressed in *S. cerevisiae* and purified as previously described (Kumar *et al*. 2014). Msh2-msh6(3MBD) was overexpressed in *E. coli* and purified as previously described (Brown *et al*. 2016).

### DNA Substrates

DNA substrates were constructed with synthetic oligonucleotides (**Table 1**). One oligonucleotide in each substrate was biotinylated for attachment to streptavidin tips (see below). Oligonucleotides were mixed at equimolar concentrations in 100 mM NaCl, 10 mM MgCl_2_, 0.1 mM EDTA, heated to 95°C for 5 minutes and allowed to cool slowly to room temperature.

### Biolayer Interferometry (BLI)

Biolayer Interferometry experiments were performed in Binding Buffer (25mM Tris-HCl pH 7.5, 1mM EDTA,100mM NaCl, 0.1mg/mL BSA, 1mM DTT, and 0.05% Tween-20) using the Octet Red96e system (ForteBio). Biotinylated DNA substrates (7.5nM) were immobilized to pre-hydrated streptavidin biosensors. Biosensors were then incubated with 50uM biocytin in Superblock buffer (ThermoFisher) to quench unbound streptavidin and equilibrated in Binding Buffer. Biosensors were incubated with Msh2-Msh3 or Msh2-msh6(3MBD) (250 nM, 125nM, 62.5nM, 31.25nM, 15.6nM, and 7.8nM) for 180 seconds. Biosensors were then moved to wells containing experiment buffer to allow for dissociation for 180 seconds. Data were collected using Data Acquisition 12.0 software (ForteBio) and analyzed with Data Analysis HT 12.0 software (ForteBio). Data were fit to 1:1 model to obtain binding kinetics. Experiments were performed in triplicate with protein obtained from at least 2 independent purifications.

### Strains and Media

All yeast transformations were performed using the lithium acetate method (Gietz *et al*. 1992). Yeast strains were derived from the S288c background; FY23 for slippage assays, FY86 for TNR expansion assays (Winston *et al*. 1995). *msh3Δ* and *msh6Δ* were constructed by amplifying a chromosomal mutant::KANMX fragment from the yeast deletion collection that was integrated into the respective chromosomal location. *msh6(3MBD)* was integrated into the endogenous MSH6 locus in a *msh3Δ* strain using pRDK4576 as described previously (Shell *et al*. 2007).

TNR substrates were integrated into all strains as described previously (Dixon *et al*. 2004; Williams and Surtees 2018): 1) (CAG)_25_ (pBL70), 2) (CTG)_25_ (pBL69), 3) scrambled (C,A,G)_25_ (pBL139) and 4) scrambled (C,T,G)_25_ (pBL138) (Dixon *et al*. 2004). The repeat tracts were in the promoter of the *URA3* reporter gene; expansions of ≥ 4 repeats alter the transcriptional start site, making the cells resistant to 5-FOA.

The microsatellite instability construct has been described previously (Sia *et al*. 1997). Repeat tracts of 1 nt. [(G)_18_], 2 nt. [(GT)_16.5_], or 4 nt. [(CAGT)_16_] were placed in-frame upstream of *URA3*. Unrepaired slippage events shift *URA3* out of frame, resulting in 5-FOA resistance.

### Microsatellite instability assay

Microsatellite instability assays were performed as described (Sia *et al*. 1997). Single colonies were grown on SC-tryptophan (SC-trp) to maintain the reporter plasmids. Colonies of ∼2 mm were selected from ≥3 independent isolates of each genetic background. Individual colonies were resuspended in 3 mL of liquid SC-trp and incubated with shaking for 20 hours at 30°C. Overnight cultures were serial diluted and plated on permissive (SC-trp) and selective (SC-trp +5-FOA) plates. Plates were incubated at 30 degrees for 2-4 days. Mutation rates were calculated by method of the median (Drake 1991). 95% confidence intervals were determined using tables of confidence intervals for the median (Nair 1940; Dixon and Massey 1969). p-values were determined by Mann-Whitney rank analysis in GraphPad Prism.

### Assay for TNR Expansion

TNR expansion assays were performed as described (Williams and Surtees 2018). Briefly, single colonies were obtained on synthetic medium (SC) lacking histidine (SC-his) or lacking histidine or uracil (SC-his-ura) for *msh6(3MBD) msh3Δ*. Individual colonies were selected from ≥3 independent isolates of each genetic background were assayed. Colonies with unexpanded tracts, which were confirmed by PCR analysis, were diluted and plated on SC-his and incubated at 30°C for 3-4 days to allow expansions to occur. Several ∼2mm colonies were selected, diluted and plated onto permissive (SC-his) and selective (SC-his +5-FOA) and incubated at 30°C from 3-4 days. Colonies were counted and expansion rates calculated as described (Drake 1991). The 95% confidence intervals were determined from tables of confidence intervals for the median (Nair 1940; Dixon and Massey 1969). p-values were determined by Mann-Whitney rank analysis (Nair 1940; Dixon and Massey 1969; Drake 1991; Sia *et al*. 1997) in GraphPad Prism.

True expansions were determined as previously described (Williams and Surtees 2015; Williams and Surtees 2018; Williams *et al*. 2020), by amplifying the reporter promoter region with SO295 (AAACTCGGTTTGACGCCTCCCATG) and SO296 (AGCAACAGGACTAGGATGAGTAGC) and digesting with *SphI* to release the TNR tract. Tract mobility was assessed by electrophoresis through a 12% native polyacrylamide gel (0.5 X TBE). At least 30 independent 5-FOA resistant colonies were characterized for each tract and genotype combination.

## Results

### Msh2-Msh3 and Msh2-msh6(3MBD) exhibit specific binding to TNR structures *in vitro*

Previous studies, including our own, have indicated that Msh2-Msh3 TNR slipped strand structure binding and stabilization is required to promote TNR expansions (Owen *et al*. 2005; Panigrahi *et al*. 2005; Kantartzis *et al*. 2012; Williams and Surtees 2015). In this study, we set out to test whether the structure-specific-binding activity of Msh2-Msh3 is *sufficient* to promote TNR expansions, using the msh6(3MBD) chimeric protein (**Figure 1**). Before testing Msh2-msh6(3MBD) activities *in vivo*, we characterized its *in vitro* DNA-binding properties, using both MMR and TNR DNA substrates (**Figure 1**)(Surtees and Alani 2006). We previously demonstrated yeast Msh2-Msh3 specificity for +8 loop DNA structures, substrates for Msh2-Msh3-mediated MMR, using electrophoretic mobility assays (EMSAs) and DNA footprinting (Surtees and Alani 2006; Lee *et al*. 2007; Brown *et al*. 2016). Human Msh2-Msh3 was similarly demonstrated to have specificity for DNA loop structures (Habraken *et al*. 1996; Wilson *et al*. 1999; Owen *et al*. 2005; Owen *et al*. 2009; Lang *et al*. 2011). Here, we used Biolayer Interferometry (BLI) to characterize yeast Msh2-Msh3 and Msh2-msh6(3MBD) binding to homoduplex DNA, a (GT)_4_ loop (+8 loop) (Surtees and Alani 2006), (*CAG*)_10_ or (*CTG*)_10_ hairpin DNA structures, using biotinylated DNA structures assembled from synthetic oligonucleotides (Surtees and Alani 2006).

Msh2-Msh3 exhibited significant non-specific binding activity to homoduplex and exhibited ∼3-to-4-fold increased affinity for the +8 loop (**Table 2;** K_D_). The relative affinities are consistent with our previous estimates using EMSA, although the current K_D_’s are ∼20 lower (Surtees and Alani 2006). The association rate (**Table 2**; k_a_) was similar for both substrates; the dissociation rate with the +8 loop (**Table 2**; k_dis_) was slower. Msh2-Msh3 also bound preferentially to *(CTG)*_*10*_ or *(CAG)*_*10*_ quasi-hairpin structures with affinities ∼4 and 7-fold higher than to homoduplex, respectively. This is consistent with human Msh2-Msh3, which exhibited binding to *CAG* tracts similar to +8 loop structures (Owen *et al*. 2005), although we note that the affinities differ. The higher affinities of Msh2-Msh3 to all three DNA structures are driven primarily by decreased k_dis_, although there was also an ∼2-fold increase in k_a_ with specific DNA substrates.

**Table 2:**
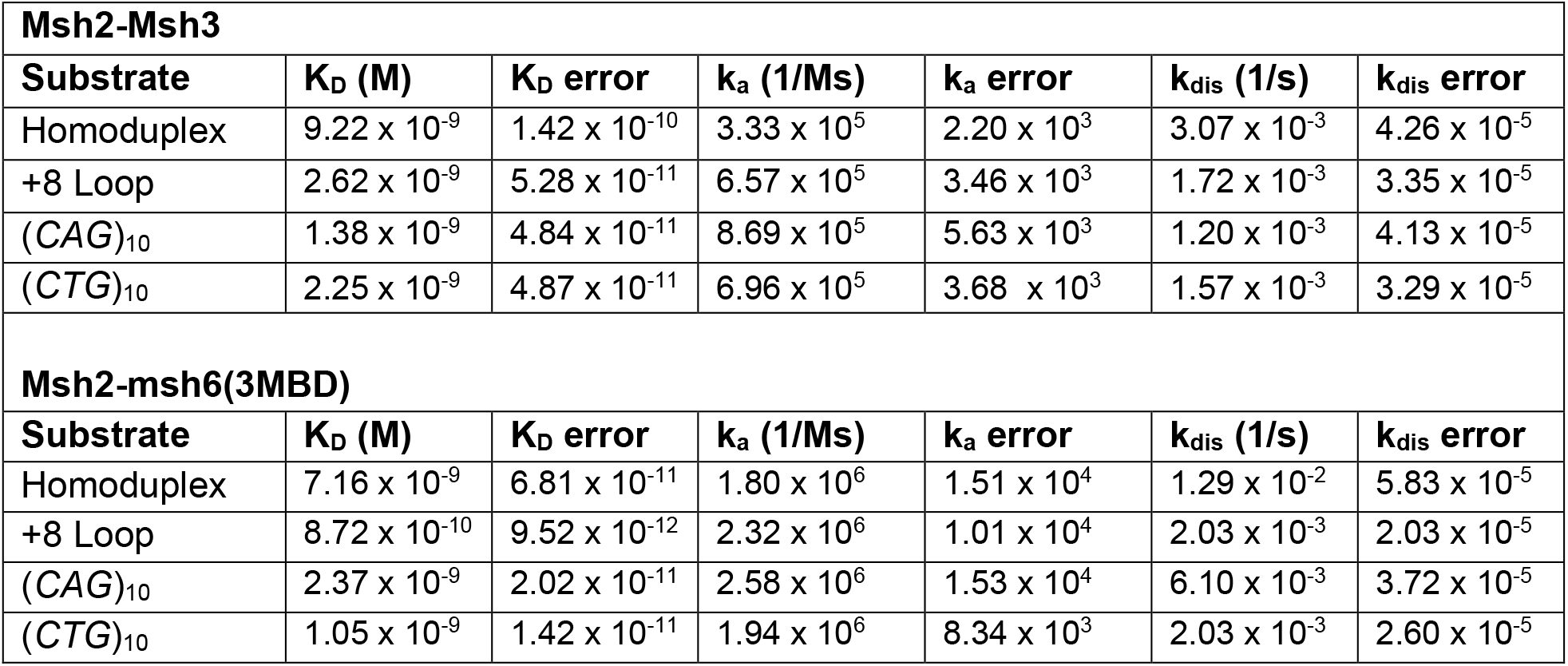
Binding kinetics of Msh2-Msh3 and Msh2-msh6(3MBD)

| Msh2-Msh3 |  |  |  |  |  |  |
| --- | --- | --- | --- | --- | --- | --- |
| Substrate | K<sub>D</sub> (M) | K<sub>D</sub> error | k<sub>a</sub> (1/Ms) | k<sub>a</sub> error | k<sub>dis</sub> (1/s) | k<sub>dis</sub> error |
| Homoduplex | 9.22 x 10<sup>-9</sup> | 1.42 x 10<sup>-10</sup> | 3.33 x 10<sup>5</sup> | 2.20 x 10<sup>3</sup> | 3.07 x 10<sup>-3</sup> | 4.26 x 10<sup>-5</sup> |
| +8 Loop | 2.62 x 10<sup>-9</sup> | 5.28 x 10<sup>-11</sup> | 6.57 x 10<sup>5</sup> | 3.46 x 10<sup>3</sup> | 1.72 x 10<sup>-3</sup> | 3.35 x 10<sup>-5</sup> |
| (CAG)<sub>10</sub> | 1.38 x 10<sup>-9</sup> | 4.84 x 10<sup>-11</sup> | 8.69 x 10<sup>5</sup> | 5.63 x 10<sup>3</sup> | 1.20 x 10<sup>-3</sup> | 4.13 x 10<sup>-5</sup> |
| (CTG)<sub>10</sub> | 2.25 x 10<sup>-9</sup> | 4.87 x 10<sup>-11</sup> | 6.96 x 10<sup>5</sup> | 3.68 x 10<sup>3</sup> | 1.57 x 10<sup>-3</sup> | 3.29 x 10<sup>-5</sup> |
| Msh2-msh6(3MBD) |  |  |  |  |  |  |
| Substrate | K<sub>D</sub> (M) | K<sub>D</sub> error | k<sub>a</sub> (1/Ms) | k<sub>a</sub> error | k<sub>dis</sub> (1/s) | k<sub>dis</sub> error |
| Homoduplex | 7.16 x 10<sup>-9</sup> | 6.81 x 10<sup>-11</sup> | 1.80 x 10<sup>6</sup> | 1.51 x 10<sup>4</sup> | 1.29 x 10<sup>-2</sup> | 5.83 x 10<sup>-5</sup> |
| +8 Loop | 8.72 x 10<sup>-10</sup> | 9.52 x 10<sup>-12</sup> | 2.32 x 10<sup>6</sup> | 1.01 x 10<sup>4</sup> | 2.03 x 10<sup>-3</sup> | 2.03 x 10<sup>-5</sup> |
| (CAG)<sub>10</sub> | 2.37 x 10<sup>-9</sup> | 2.02 x 10<sup>-11</sup> | 2.58 x 10<sup>6</sup> | 1.53 x 10<sup>4</sup> | 6.10 x 10<sup>-3</sup> | 3.72 x 10<sup>-5</sup> |
| (CTG)<sub>10</sub> | 1.05 x 10<sup>-9</sup> | 1.42 x 10<sup>-11</sup> | 1.94 x 10<sup>6</sup> | 8.34 x 10<sup>3</sup> | 2.03 x 10<sup>-3</sup> | 2.60 x 10<sup>-5</sup> |

We next tested the DNA-binding kinetics of Msh2-msh6(3MBD), which replaces the Msh6 MBD with Msh3 MBD within the context of Msh2-Ms6 (**Figure 1**) (Surtees and Alani 2006; Shell *et al*. 2007; Brown *et al*. 2016). Msh2-msh6(3MBD) displayed a higher (∼8-fold) affinity (**Table 2**; K_D_) for the +8 loop structure compared to the homoduplex DNA structure, consistent with previous observations (Shell *et al*. 2007; Brown *et al*. 2016) and with Msh2-Msh3 binding (**Table 2**) (Surtees and Alani 2006). Msh2-msh6(3MBD) also exhibited increased affinity for both TNR hairpin structures, relative to homoduplex, with affinities similar to its affinity for the +8 loop substrate (**Table 2**; K_D_), as observed with Msh2-Msh3 (**Table 2**). Msh2-msh6(3MBD) exhibited similar k_a_’s for all of the substrates, which were somewhat higher than Msh2-Msh3 k_a_’s for the same substrates. As with Msh2-Msh3, the increased affinity to specific DNA structures was largely driven by decreased dissociation rates (**Table 2**; k_dis_). These data indicate that Msh2-Msh3 and Msh2-msh6(3MBD) exhibit similar DNA binding affinities and that Msh2-msh6(3MBD) binds TNR structures with an affinity similar to its affinity for MMR (+8 loop) structures and is therefore competent to recognize these structures, should they form *in vivo*.

### *msh6(3MBD)* retains some Msh2-Msh3-specific MMR function

We integrated the *msh6(msh3MBD)* construct (Shell *et al*. 2007) into the chromosome, replacing endogenous *MSH6*, in a *msh3Δ* background. This results in a single MSH complex *in vivo*, Msh2-msh6(3MBD) (**Figure 1**). Previous work has demonstrated that *msh6(msh3MBD) msh3Δ* has *in vivo* MMR activity, indicating that the complex is functional, although both Msh2-Msh6-and Msh2-Msh3-mediated MMR were compromised to some extent (Shell *et al*. 2007). *msh6(msh3MBD) msh3Δ* function was tested with reporter assays that select for -1 frame shift deletion or +2 frame shift insertions (Shell *et al*. 2007), substrates for both Msh2-Msh3 and Msh2-Msh6 (Marsischky *et al*. 1996; Sia *et al*. 1997). *msh6(3MBD) msh3Δ* elevated mutation rates (30-70-fold increases), but not the synergistic >1000-fold increase observed in *msh3Δ msh6Δ* (Shell *et al*. 2007), indicating that *msh6(3MBD)* retained significant MMR function. Here we tested whether the chimeric protein would act in repair of larger IDLs that is dependent almost exclusively on Msh2-Msh3-mediated repair (Sia *et al*. 1997; Lee *et al*. 2007; Kumar *et al*. 2013). We used a microsatellite instability reporter plasmid that places a tetranucleotide [*CAGT*]_16_ repeat in-frame upstream of *URA3* (Sia *et al*. 1997). Unrepaired DNA slippage events alters the *URA3* reading frame, resulting in 5-FOA resistance, which allows selection of these slippage events. Previous work demonstrated that these slippage events result primarily in deletions (Sia *et al*. 1997; Lamb *et al*. 2022). Compared to wild-type, *msh3Δ* increased the 4 nt. slippage rate by 58-fold; *msh6Δ*, exhibited only a 3-fold increase, consistent with its limited role in repair of these longer in/dels (**Figure 2**; blue circles). *msh2Δ* exhibited a slippage rate similar to *msh3Δ* with this 4 nt. repeat (57- and 62-fold increases over wild-type, respectively) (Sia *et al*. 1997; Lee *et al*. 2007), consistent with *MSH6* playing only a minor role in repair of larger IDL repair. *msh6(3MBD) msh3Δ* partially complemented the *msh3Δ*, with an intermediate slippage rate, with a 24-fold increase (**Figure 2**). This incomplete complementation is consistent with distinct repair mechanisms for Msh2-Msh3 versus Msh2-Msh6, but is consistent with *msh6(3MBD*) allowing recognition of IDL substrates *in vivo*. We also tested *msh6(3MBD) msh3Δ* function in repair of 1 nt ([G]_18_) and 2 nt ([GT]_16.5_) repeats (Sia *et al*. 1997; Lee *et al*. 2007) *msh6(3MBD) msh3Δ* partially complemented repair of 1 nt. slippage events, compared to *msh3Δ*, but not repair of 2 nt. slippage events. We note that *msh2Δ* exhibited synergistic slippage rates with 1 nt. and 2 nt. repeat reporters (Sia *et al*. 1997; Lee *et al*. 2007), consistent with partial function of *msh6(3MBD)* in repair of these repeats (**Figure 2**) (Shell *et al*. 2007).

**Figure 2.**
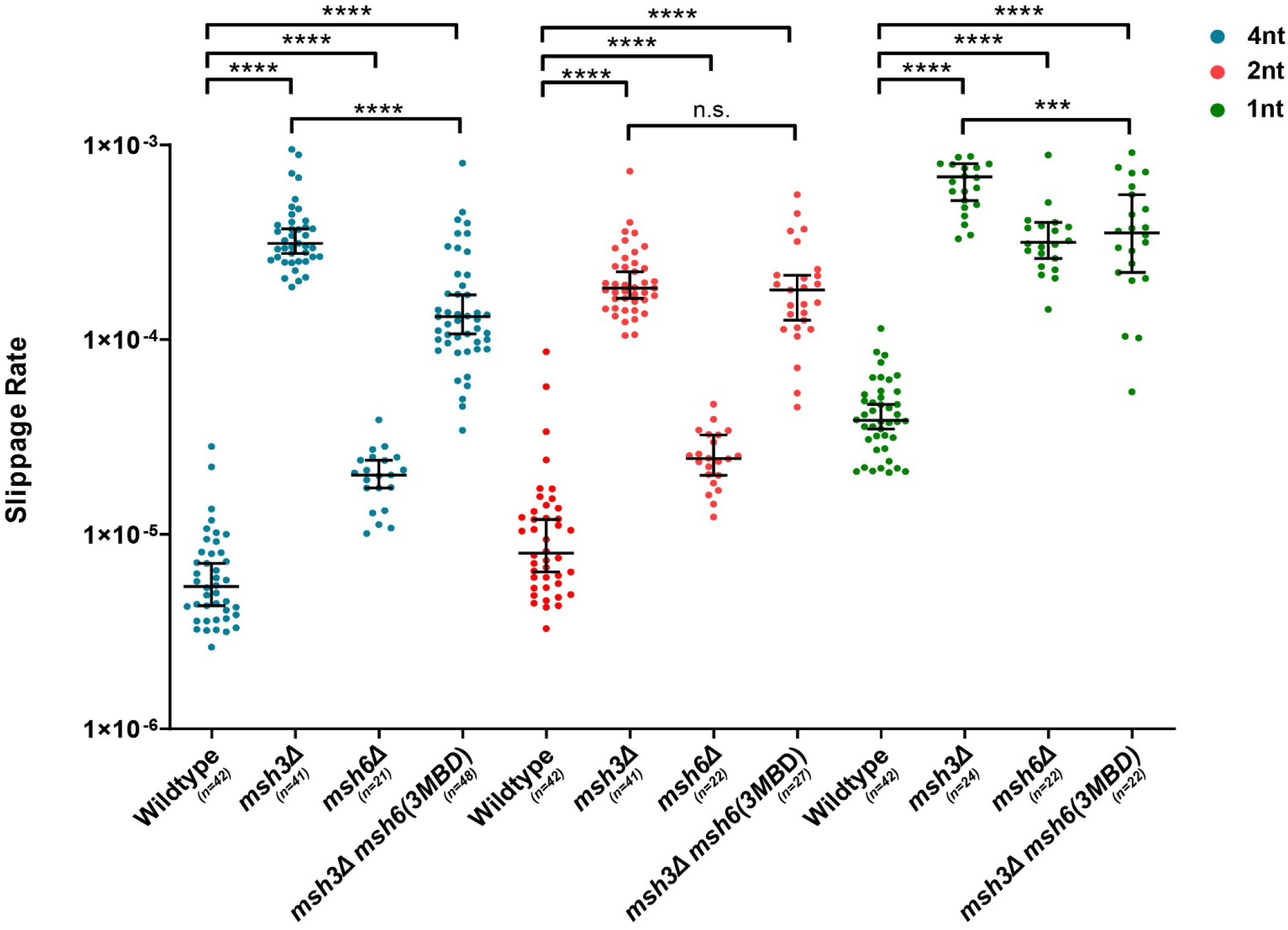
Mutation rate in a slippage mutation rate assay. The rate of slippage events, which push *URA3* out of frame and allow selection in the presence of 5-FOA, was determined in different genetic backgrounds, as indicated. Slippage rate in the presence of mononucleotide (green) dinucleotide (green) and tetranucleotide (blue) repeats were determined. Median rates *[95% confidence interval]* are as follows: **<u>4nt (blue):</u> wild-type**: 5.38 × 10^−6^ *[4.30 × 10*^*-6*^ *– 7.08 × 10*^*-6*^], ***msh3Δ***: 3.12 × 10^−4^ *[2.77 × 10*^*-4*^ *– 3.71 × 10*^*-4*^], ***msh6Δ***: 2.01 × 10^−5^ *[1.73 × 10*^*-5*^ *– 2.4 × 10*^*-5*^], ***msh6(3MBD) msh3Δ***: 1.29 × 10^−4^ *[1.07 × 10*^*-4*^ *– 1.70 × 10*^*-4*^], **<u>2nt (red):</u> wild-type**: 7.99 × 10^−6^ *[6.14 × 10*^*-6*^ *– 1.20 × 10*^*-5*^], ***msh3Δ***: 1.84 × 10^−4^ *[1.63 × 10*^*-4*^*-2.23 × 10*^*-4*^], ***msh6Δ***: 2.45 × 10^−5^ *[2.01 × 10*^*-5*^ *– 3.24 × 10*^*-5*^], ***msh6(3MBD) msh3Δ***: 1.80 × 10^−4^ *[1.26 × 10*^*-4*^ *– 2.14 × 10*^*-4*^], **<u>1nt (green):</u> wild-type**: 3.84 × 10-5 *[3.47 × 10*^*-5*^ *– 4.64 × 10*^*-5*^], ***msh3Δ*** = 6.85 × 10^−4^ *[5.18 × 10*^*-4*^ *– 8.02 × 10*^*-4*^], ***msh6Δ*** = 3.17 × 10^−*4*^ *[2.61 × 10*^*-4*^ *– 4.00 × 10*^*-4*^], ***msh6(3MBD) msh3Δ*** = 3.53 × 10^−4^ *[2.21 × 10*^*-4*^ *– 5.55 × 10*^*-4*^]. Error bars indicate 95% confidence intervals. p-values were calculated using Mann-Whitney test, ****: p < 0.0001, ***: 0.0001< p < 0.001.

### *msh6(3MBD)* does not promote TNR expansions *in vivo*

We next sought to determine whether the Msh3 MBD was sufficient to promote expansions *in vivo* (Shell *et al*. 2007). We used an *in vivo* TNR expansion assay in *msh6(3MBD) msh3Δ* to determine the effect of this construct on *CAG* and *CTG* expansion rates. This assay, described previously (Miret *et al*. 1998; Williams *et al*. 2020), places a *URA3* reporter gene downstream of a promoter that encodes a (*CNG*)_25_ repeat tract. If an expansion of 4 or more repeats occurs in the tract (≥29 repeats), *URA3* will not be transcribed, leading to 5-FOA resistance (Williams and Surtees 2018). We previously demonstrated that *msh3Δ* reduces the expansion rate for (*CAG)*_*25*_ and (*CTG)*_*25*_ repeat tracts 5-fold and 30-fold, respectively, while the expansion rate was slightly increased in *msh6Δ* (**Figure 3**) (Kantartzis *et al*. 2012). In *msh6(msh3MBD) msh3Δ*, the expansion rate for the (*CAG)*_*25*_ tract was very similar to that of the *msh3Δ*, while the *(CTG)*_*25*_ expansion rate was slightly lower than *msh3Δ* (**Figure 3**). These results indicate that *msh6(3MBD)* does not complement a *msh3Δ*. Therefore, the Msh3 MBD is not sufficient to promote expansions and does not confer the capacity to promote TNR expansions on Msh6, despite the fact that Msh2-msh6(msh3MBD) is able to bind specifically to TNR structures.

**Figure 3.**
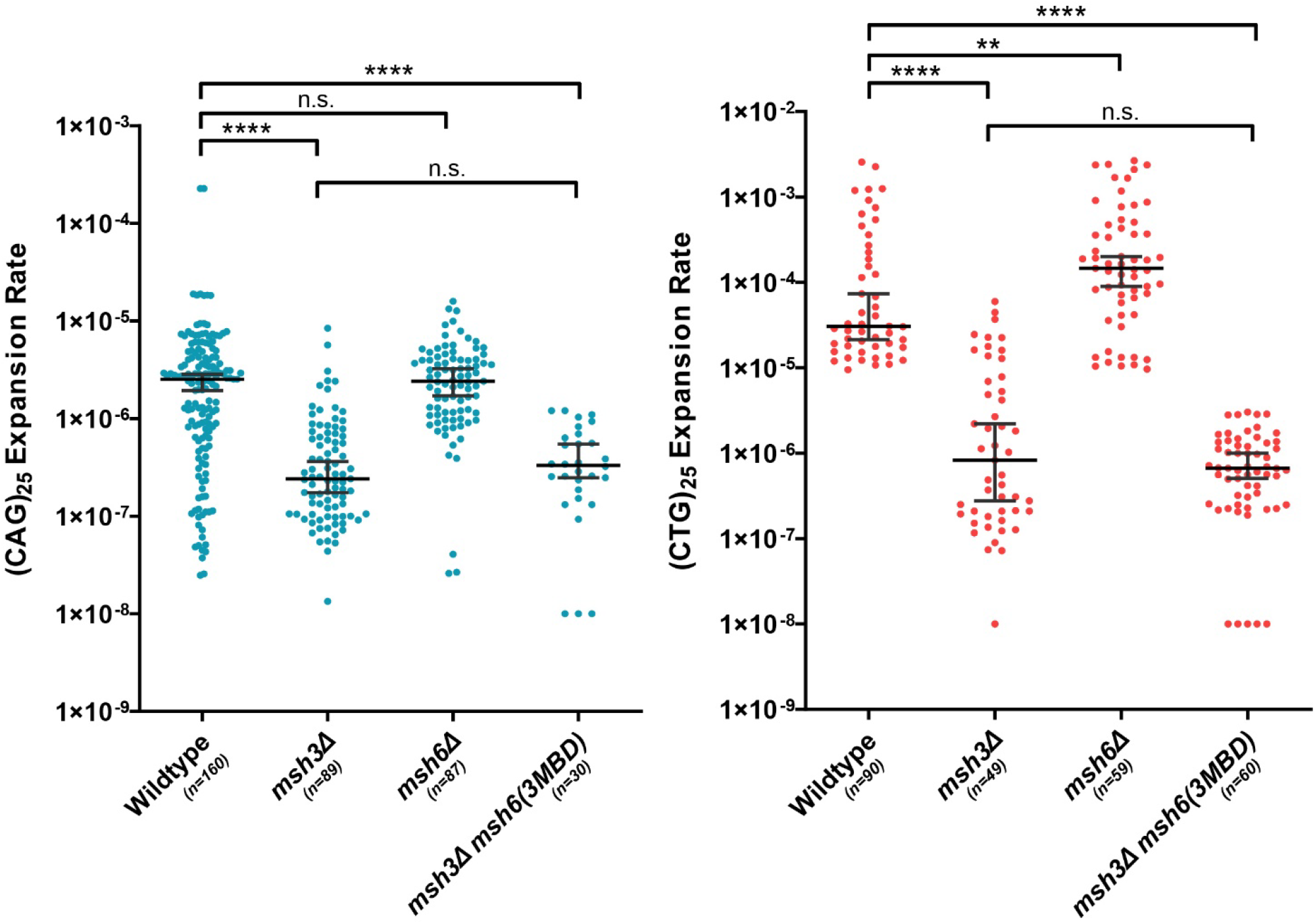
TNR expansion rates *in vivo*. The rate of expansion events, which prevent expression of *URA3* and allow selection in the presence of 5-FOA, was determined in different genetic backgrounds, as indicated. Expansion rates of *(CAG)*_*25*_ tracts (blue) and *(CTG)*_25_ tracts (red) were measured. Error bars indicate 95% confidence intervals. Median rates *[95% confidence interval]* are as follows: **(*<u>CAG</u>*)**_**<u>25</u>**_ **<u>(blue)</u>**: **wild-type**: 1.1 × 10^−6^ *[7.0 × 10*^*-7*^ *-1.9 × 10*^*-6*^], ***msh3Δ***: 2.4 × 10^−7^ *[1.6 × 10*^*-7*^ *-2.8 × 10*^*-7*^], ***msh6Δ***: 2.4 × 10^−6^ *[1.7 × 10*^*-6*^ *-2.9 × 10*^*-6*^], ***msh6(3MBD) msh3Δ***: 2.5 × 10^−7^ *[1.3 × 10*^*-7*^ *-3.4 × 10*^*-7*^]; **(*<u>CTG</u>*)**_**<u>25</u>**_ **(red): wild-type**: 2.2 × 10^−5^ *[1.7 × 10*^*-5*^ *-2.9 × 10*^*-5*^], ***msh3Δ***: 1.0 × 10^−6^ *[2.8 × 10*^*-7*^ *-2.2 × 10*^*-6*^], ***msh6Δ***:1.5 × 10^−4^ *[9.0 × 10*^*-5*^ *-2.0 × 10*^*-4*^], ***msh6(3MBD*) *msh3Δ***: 6.6 × 10^−7^ *[5.0 × 10*^*-7*^ *-9.8 × 10*^*-7*^]. Scrambled (C,A,G)_25_ and (C,T,G)_25_ tracts were also tested and median rates are as follows: **<u>(C,A,G)</u>**_**<u>25</u>**_ **wild-type** (n = 180) <1.0 × 10^−8^, ***msh3Δ*** (n = 90) <1.0 × 10^−8^, ***msh6Δ*** (n = 30) <1.0 × 10^−8^, ***msh6(3MBD) msh3Δ*** (n = 20) = 2.1 × 10^−7^ *[1.6 × 10*^*-7*^ *-3.5 × 10*^*-7*^], **<u>(C,T,G)</u>**_**<u>25</u>:**_ **wild-type** (n = 90) <1.0 × 10^−8^, ***msh3Δ*** (n = 90) <1.0 × 10^−8^, ***msh6Δ*** (n = 51) <1.0 × 10^−8^, ***msh6(3MBD) msh3Δ*** (n = 20) = 2.5 × 10^−7^ *[1.5 × 10*^*-7*^ *-3.6 × 10*^*-7*^]. p-values were calculated using Mann-Whitney test, **: 0.001 < p < 0.01, ****: p < 0.0001. Please note the scale on the y-axis for *CTG* expansions is extended.

Mutations in *URA3* can also lead to 5-FOA resistance. Therefore, we determined the proportion of true expansions by amplifying the TNR tract from 5-FOA-resistant colonies and determining tract lengths by gel electrophoresis (**Figure 4**) (Williams *et al*. 2020). In wild-type cells, 90% of *CAG* and >99% of *CTG* tracts exhibited true expansions. In *msh3Δ*, 63% of *CAG* and 91% of *CTG* tracts were *bona fide* (Kantartzis *et al*. 2012). In *msh6(3MBD) msh3Δ*, 7% of *CAG* tracts and 33% of *CTG* tracts exhibited true expansions (**Table 3**). The remaining tracts were either stable in length or exhibited tract contractions. We also observed a high rate of 5-FOA resistance in these strains with scrambled tracts, which do not expand (**Figure 3 legend**), indicating that, in *msh6(3MBD) msh3Δ*, which has a high mutation rate (**Figure 2**), mutations other than TNR tract expansions are leading to 5-FOA resistance. This may be caused by mutations in *URA3* or *URA6* (Armstrong *et al*. 2024). Therefore, the calculated expansion rates for *msh6(3MBD) msh3Δ* are overestimates; a corrected rate for *msh6(3MBD) msh3Δ* TNR expansions would go from 2.5 × 10^−7^ to 1.8 × 10^−8^ (*CAG*) or from 6.6 × 10^−7^ to 2.2 × 10^−7^ (*CTG*). Thus, measuring TNR expansion rates with this assay becomes more complicated as the background mutation rate increases for a given genotype, increasing the probability of observing 5-FOA resistance without expansion. Nonetheless, our data suggest that the *CAG* and *CTG* expansion rates for the chimeric complex are ∼8-fold and ∼4-fold lower than in the absence of *MSH3*, respectively, with corrected expansion rates of 1.5 × 10^−7^ (*msh3Δ CAG*) and 9.7 × 10^−7^ (*msh3Δ CTG*).

**Table 3:**
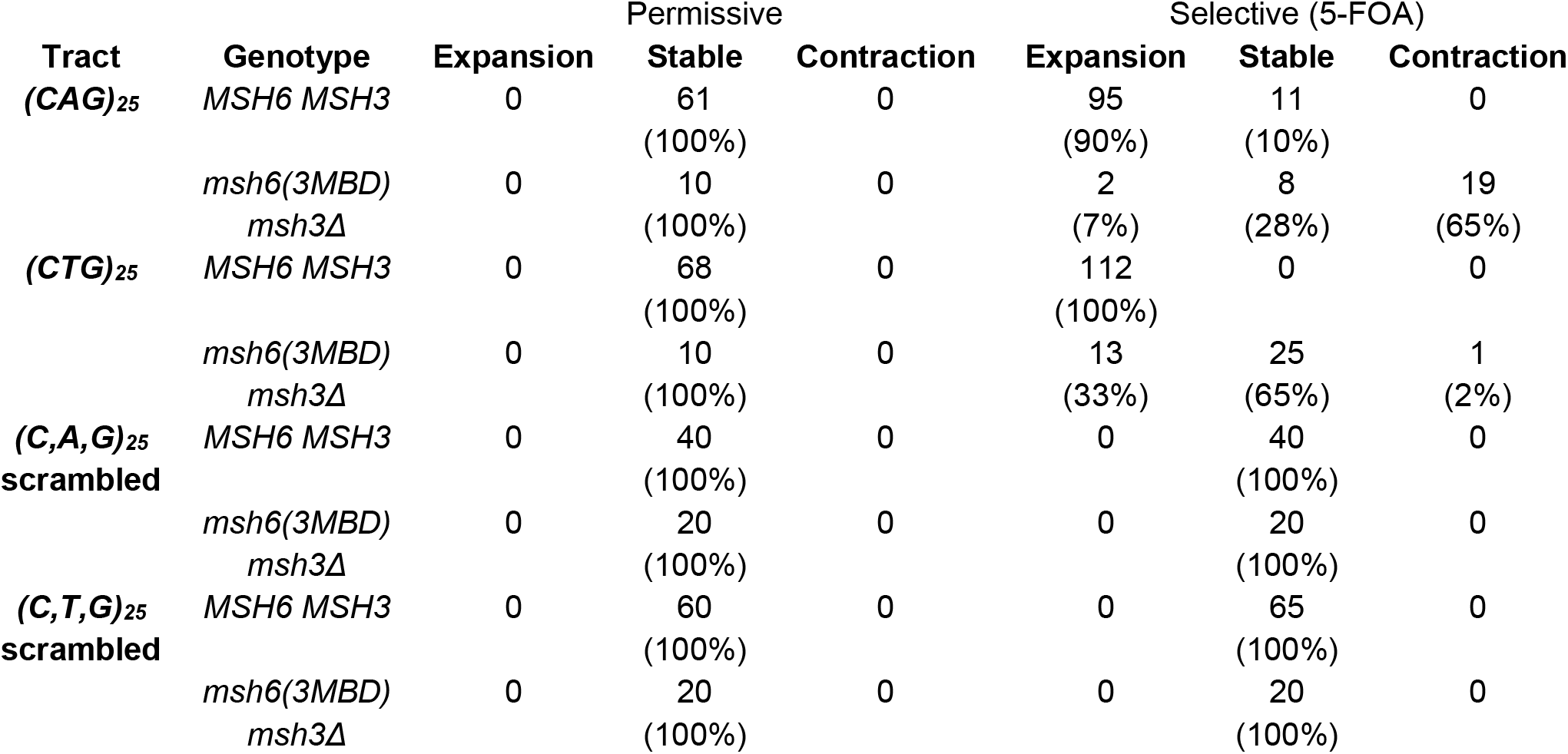
Proportion of true expansions in 5-FOA resistant colonies.

**Figure 4.**
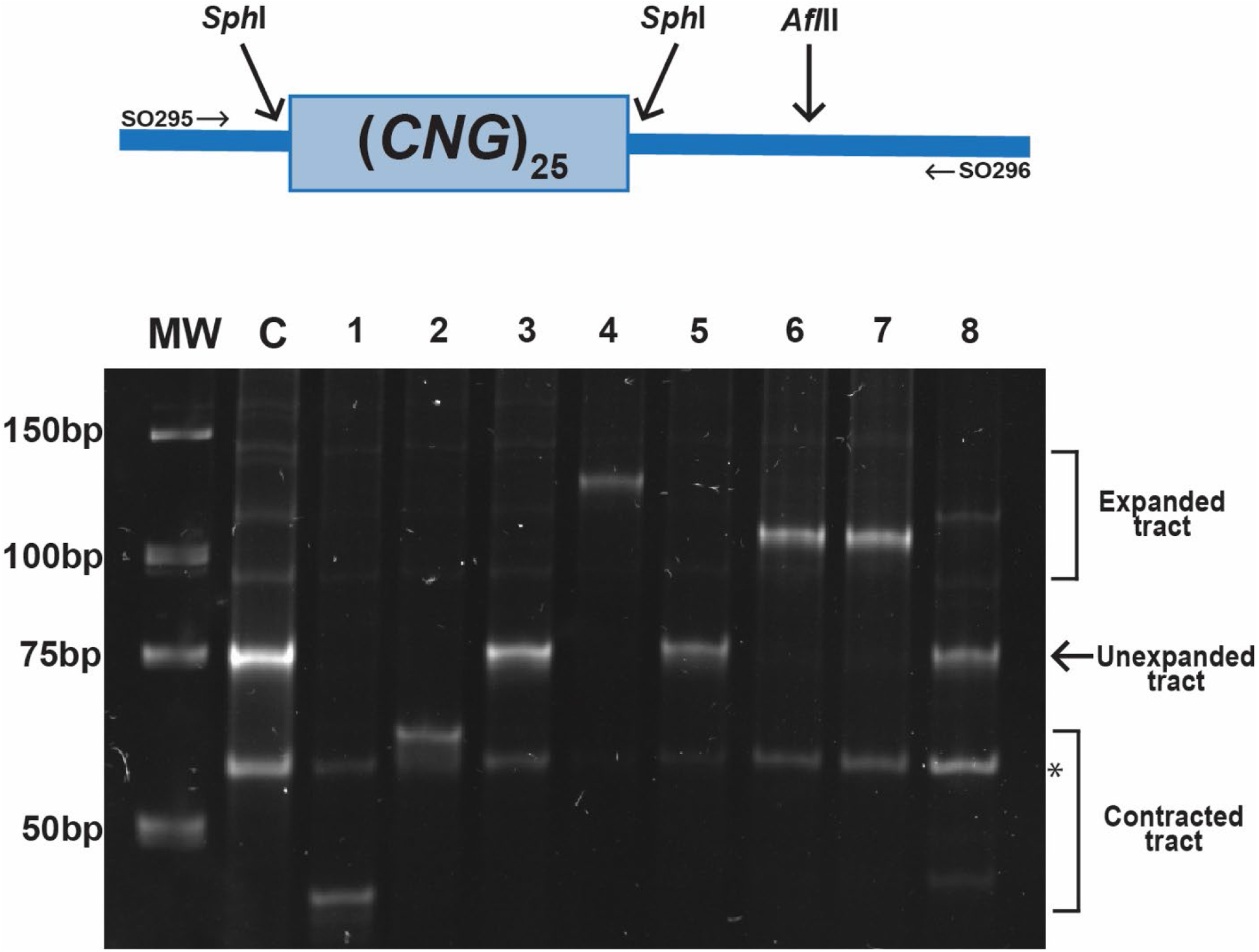
TNR tract length in reporters recovered from 5-FOA resistant colonies. *Top:* Schematic of PCR product used to measure tract lengths of 5-FOA resistant colonies. Amplification with SO295 and SO296 results in a 188bp product. Digestion with *Sph*I cuts on either side of the tract, releasing the 75bp tract. The remaining 73bp product adjacent to the tract is digested with *Afl*II, allowing for visualization of the TNR tract. *Bottom:* Representative gel of digested TNR tracts amplified from 5-FOA resistant colonies from wild-type and *msh6(3MBD) msh3Δ*. These tracts include contracted (lanes 1 and 2), expanded (lanes 4, 6, and 7) and stable TNR tracts (lanes 3, 5, and 8). * denotes band that results from *Afl*II digestion.

## Discussion

We demonstrated that, *in vitro*, Msh2-msh6(3MBD) exhibited DNA structure binding affinities for loop and TNR structures that were comparable to Msh2-Msh3, indicating that Msh2-msh6(3MBD) is able to recognize and bind TNR structures *in vivo* (**Table 1**). We demonstrated that *msh6(3MBD) msh3Δ* is sufficient to allow some repair of a Msh2-Msh3-specific DNA errors (in/dels), partially complementing the loss of *MSH3* in 4 nt. loop repair *in vivo* (**Figure 2**). We hypothesize that differences in communication between the DNA-binding and ATPase domains of Msh2-Msh3 versus Msh2-Msh6 are reflected in this partial complementation. In contrast, this level of activity in *msh6(3MBD) msh3Δ* is not sufficient to promote MMR-mediated *CAG/CTG* expansions (**Figure 3**). In fact, the true TNR expansion rates appear to be lower in *msh6(3MBD) msh3Δ* than in *msh3Δ*. Thus, we propose that distinct Msh3-specific molecular requirements beyond Msh3 MBD are necessary for promoting TNR expansions, including DNA-mediated modulation of Msh2-Msh3 ATP-binding and hydrolysis and interactions with MLH complexes. Further, Msh2-msh6(3MBD), an MSH complex that is able to bind MMR and TNR structures (**Table 2**) but not coordinate efficient repair (**Figure 2,3**), appears to block background TNR expansions.

Msh2-Msh3 DNA-binding to distinct structures is communicated to the ATPase domain through the connector domain, modulating its ATP binding, hydrolysis and turnover activities to promote repair (Owen *et al*. 2005; Owen *et al*. 2009; Gupta *et al*. 2011; Lang *et al*. 2011; Kumar *et al*. 2014). ATP promotes Msh2-Msh3 dissociation from the DNA, promoting recycling of Msh2-Msh3 (Surtees and Alani 2006; Brown *et al*. 2016). Different DNA structures modulate Msh2-Msh3 ATP binding and hydrolysis (Owen *et al*. 2005; Surtees and Alani 2006; Owen *et al*. 2009; Kumar *et al*. 2014).

Critically, the Msh2-Msh3 ATP-binding domains are distinct from those of Msh2-Msh6, with regulated access to the Msh3 nucleotide binding pocket (Gupta *et al*. 2011; Kumar *et al*. 2013). Altered regulation of ATP binding and/or hydrolysis disrupts Msh2-Msh3-mediated MMR, but not Msh2-Msh3’s DSBR activity (Kumar *et al*. 2013). Therefore, Msh2-msh6(3MBD) may misregulate the ATPase domain through incorrect signal transduction after DNA binding and altered access to the nucleotide binding pocket. Our previous data indicate that Msh2-msh6(3MBD) has a higher ATPase activity than Msh2-Msh3, more similar to Msh2-Msh6, while being stimulated by Msh2-Msh3-specific DNA substrates (Brown *et al*. 2016). One possibility is that this elevated ATPase activity increases turnover of Msh2-msh6(3MBD), impairing both Msh2-Msh3-mediated MMR and TNR expansions, although apparently not to the same extent (**Figure 2**) (Shell *et al*. 2007). This may be a result of distinct DNA structure-specific allosteric changes within the Msh complex. We propose that increased turnover precludes the Msh complex from targeting the DNA structures for either MMR or TNR expansion, leading to defects in both Msh2-Msh3-mediated pathways. Together, our data indicates that Msh2-Msh3’s MMR activity that is specifically required for promoting TNR expansions.

ATP-induced conformational changes are required for Msh2-Msh3’s interaction with MLH complexes and stimulates their endonuclease activity. MutLα (yeast Mlh1-Pms1) is the primary Mlh complex in MMR, although MutLγ (yeast Mlh1-Mlh3) also plays a minor, largely Msh2-Msh3-specific, role in MMR. Both MutLα and MutLγ promote TNR expansions *in vivo* in mammalian systems in a Msh2-Msh3-specific manner (Pinto *et al*. 2013; Zhao *et al*. 2018; Hayward *et al*. 2020; Kadyrova *et al*. 2020; Miller *et al*. 2020; Roy *et al*. 2021; Lee *et al*. 2022), likely by nicking and promoting excision of the template leading to expansions (Pluciennik *et al*. 2013; Kadyrova *et al*. 2020). Our data support the hypothesis that this is not simply a result of altered DNA-binding specificity, but rather that Msh2-Msh3-specific Mlh interactions and activation are required for TNR expansions and are missing in Msh2-msh6(3MBD). We note that human Msh2-Msh3 interactions with MutLα are mediated through the PCNA interaction motif (PIP box) with the Msh3 N-terminal region (Pluciennik *et al*. 2013). In contrast, the Msh6 NTR or PIP box is not required for this interaction (Iyer *et al*. 2008), therefore any Msh2-msh6(3MBD) interactions with MLH complexes are expected to be quite different from Msh2-Msh3-MLH interactions. We propose that Msh2-Msh3-mediated TNR expansions require the Msh2-Msh3-mediated MMR pathway to be fully functional and intact. This would include proper DNA binding and appropriate signal transduction for regulation of ATP binding/hydrolysis and subsequent Mlh interactions. This is consistent with Msh2-Msh3 playing an active, pathogenic role in promoting TNR expansions.

## Data Availability Statement

All strains and plasmids are available upon request.

## Funding

This work was supported by American Cancer Society Research Scholar Grant (RSG-14-2350-01), Pfizer and the National Science Foundation (MCB grant #2325415) to JAS.

## Acknowledgments

We thank Dr. Natalie Lamb and Brett Irwin and other members of the Surtees lab for technical assistance and for helpful discussions. We are grateful to Dr. Mark Sutton for constructive comments on the study and the manuscript.

## Conflict of Interest

The authors declare they have no conflict of interest.

## Notes

### Competing Interest Statement

The authors have declared no competing interest.

